# Structural basis for heme-dependent NCoR binding to the transcriptional repressor REV-ERBβ

**DOI:** 10.1101/2020.05.05.079277

**Authors:** Sarah A. Mosure, Jinsai Shang, Paola Munoz-Tello, Douglas J. Kojetin

## Abstract

Heme is the endogenous ligand for the constitutively repressive REV-ERB nuclear receptors, REV-ERBα (NR1D1) and REV-ERBβ (NR1D2), but how heme regulates REV-ERB activity remains unclear. While cellular studies indicate heme is required for the REV-ERBs to bind the corepressor NCoR and repress transcription, fluorescence-based biochemical assays and crystal structures suggest that heme displaces NCoR. Here, we show that heme artifactually influences detection of NCoR interaction in fluorescence-based assays. However, using fluorescence-independent methods, isothermal titration calorimetry and NMR spectroscopy, we demonstrate that heme directly increases REV-ERBβ ligand-binding domain (LBD) binding affinity for NCoR. We further report two crystal structures of REV-ERBβ LBD cobound to heme and NCoR peptides, which reveal the structural basis for heme-dependent NCoR binding to REV-ERBβ. By resolving previous contradictory biochemical, structural, and cellular studies, our findings should facilitate renewed progress toward understanding heme-dependent REV-ERB activity.

## INTRODUCTION

Nuclear receptors (NRs) are a superfamily of transcription factors that evolved to bind endogenous small molecule ligands (*1*). Defining the molecular basis for NR regulation by their natural ligands provides important insight into how extracellular and intracellular signals are transmitted into changes in gene expression. This information helps identify processes that may be dysregulated in disease and informs the design of synthetic NR ligands with therapeutic potential.

The REV-ERBs (REV-ERBα/NR1D1 and REV-ERBβ/NR1D2) are two closely related NRs with critical roles in mammalian physiology, including maintenance of the circadian rhythm, metabolic processes, and immune function (*2*). The REV-ERBs are unique among NRs because they lack the C-terminal Activation Function-2 (AF-2) helix 12 within their ligand-binding domain (LBD) important for binding transcriptional coactivator proteins, suggesting they should interact exclusively with transcriptional corepressor proteins and solely repress transcription (*3, 4*), which is supported by evidence that the REV-ERBs constitutively repress target genes (*5*).

The unique structure and repressive activity of the REV-ERBs raises a compelling dilemma. For other NRs that have an AF-2 helix 12 and interact with both transcriptional coactivator and corepressor proteins, ligand binding is important for dictating coregulator preference and determining the activation or repressive effect on target gene transcription (*6*). However, if the REV-ERBs are structurally primed to interact exclusively with corepressors, the role of ligands in regulating REV-ERB activity is unclear. One possibility is that without ligand, REV-ERBs do not interact with corepressors and are transcriptionally inactive—a hypothesis supported by cell-based evidence. The iron-centered porphyrin heme was identified as an endogenous physiological REV-ERB ligand (*7, 8*). Cellular studies demonstrated that heme is required for the REV-ERBs to interact with the transcriptional corepressor protein Nuclear receptor CoRepressor-1 (NCoR). Disruption of heme binding by point mutation or chemical depletion of cellular heme inhibits the interaction between the REV-ERBs and NCoR and repression of target genes (*7-9*).

Structural data would help establish that heme directly produces a REV-ERB LBD conformation that enhances NCoR binding. However, published biochemical and structural studies contradict cell-based evidence (*7-11*). In fluorescence-based biochemical assays, heme was shown to dose-dependently displace NCoR interaction domain (ID) peptides from the REV-ERBs (*7-11*), and comparison of REV-ERB LBD crystal structures bound either to heme alone or an NCoR ID peptide alone in the absence of heme indicated that the heme-bound conformation would directly clash with the NCoR-bound conformation (*10, 11*) (**Fig. 1**). Due to the inherent complexity within cells, observations from cell-based assays are not guaranteed to be directly attributable to components tested, while biochemical and structural studies using the well-defined, purified components establish which effects are direct. Thus, the conflicting biochemical, structural, and cellular data have cast doubt on whether heme directly promotes REV-ERB interaction with NCoR and generally prevented progress toward understanding the molecular basis for REV-ERB activity.

**Fig. 1.**
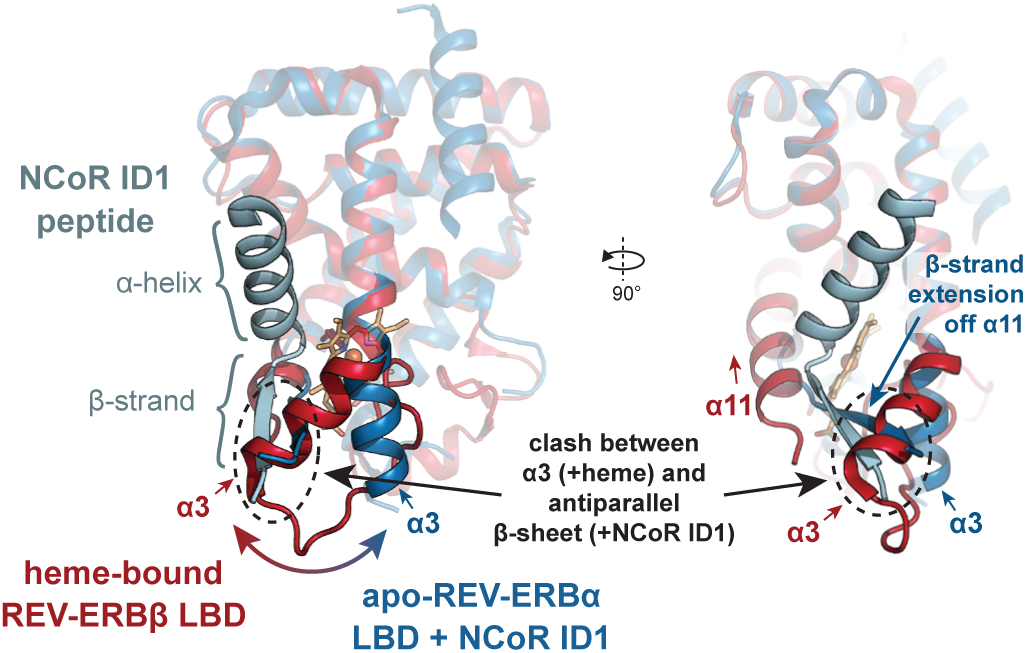
Structural clash between heme- or NCoR-bound REV-ERB LBD conformations. Crystal structures of heme-bound REV-ERBβ LBD (dark red cartoon with heme shown as orange sticks; PDB 3CQV) and apo-REV-ERBα LBD bound to NCoR ID1 peptide (dark blue cartoon with ID1 shown as light blue cartoon; PDB 3N00). Dashed black oval indicates the clash between the helix 3 (α3) conformation in the heme-bound structure and the beta-sheet formed between the NCoR ID1 peptide and the C-terminal extension of helix 11 (α11) in the NCoR ID1-bound structure.

Here, we provide evidence that heme-dependent fluorescence assay artifacts have led to an inaccurate conclusion that heme physically displaces NCoR from the REV-ERBs. Using two fluorescence-independent methods, isothermal titration calorimetry (ITC) and NMR spectroscopy, we show heme directly increases REV-ERBβ LBD affinity for an NCoR ID peptide and the entire NCoR Receptor Interaction Domain (RID), which is further supported by two crystal structures that we solved of REV-ERBβ LBD cobound to heme and NCoR ID peptides. Importantly, our work here provides biochemical and structural data that cohesively support the hypothesis that the REV-ERBs require heme to interact with corepressors in cells to actively repress transcription.

## RESULTS

### Heme-dependent artifacts in fluorescence-based assays

The NCoR corepressor interacts with the REV-ERB LBD through ID peptide motifs present within the NCoR RID (*12*) (**Fig. 2A**). Previous studies using purified recombinant REV-ERBα and REV-ERBβ LBD showed that heme reduced LBD affinity for and dose-dependently displaced peptides derived from NCoR ID motifs peptides in fluorescence-based assays (*7-11*). We observed that heme had a similar effect on REV-ERBβ LBD interaction with FITC-labeled NCoR ID peptides in time-resolved fluorescence energy transfer (TR-FRET) experiments (**Fig. 2B**). While heme itself does not fluoresce, it absorbs broadly across the UV-visible spectrum and heme-bound REV-ERBβ LBD also shows nonzero absorbance through wavelengths up to at least 700 nm (**Fig. 2C**). These absorption properties indicate heme is capable of quenching fluorescence. Indeed, several studies have documented heme-dependent fluorescence quenching while exploring heme biology. For fluorescence-based cellular heme reporters, the presence of labile heme is identified when it binds to a heme-binding protein and quenches the fluorescence of a tandemly linked fluorescent protein (*13*). Heme quenching of pocket tryptophan and tyrosine residue fluorescence is known to limit the use of tryptophan or tyrosine fluorescence quenching assays to study ligand cobinding to heme-bound proteins (*14*). An additional study warned against the use of fluorescence-based substrates in assays studying heme-binding proteins (e.g., catalase, hemoglobin) because heme quenching of the fluorescent signal causes assay artifacts (*15*). Consistent with these studies, we found that heme dose-dependently quenched the fluorescence of fluorophores with excitation and emission wavelengths ranging from 495 nm to 785 nm (**Fig. 2D**). Although the quenching effect was more pronounced for fluorophores with excitation/emission wavelengths near the heme maximal absorbance (∼400 nm; FITC and PE), heme also quenched the far-red fluorophores (eFluor780 and APC) traditionally thought to be insensitive to heme (*13*). These data indicate that fluorescence is highly sensitive to heme-dependent quenching effects and assays in which heme is near the fluorophore (i.e., in a cobinding scenario with a fluorescently-labeled peptide) may be particularly susceptible to artifacts. Importantly, these findings suggest that the decrease in signal in fluorescence-based NCoR ID peptide binding assays could be attributable to heme quenching the fluorescent signal itself instead of heme physically displacing ID peptides from the LBD (**Fig. 2E**).

**Fig. 2.**
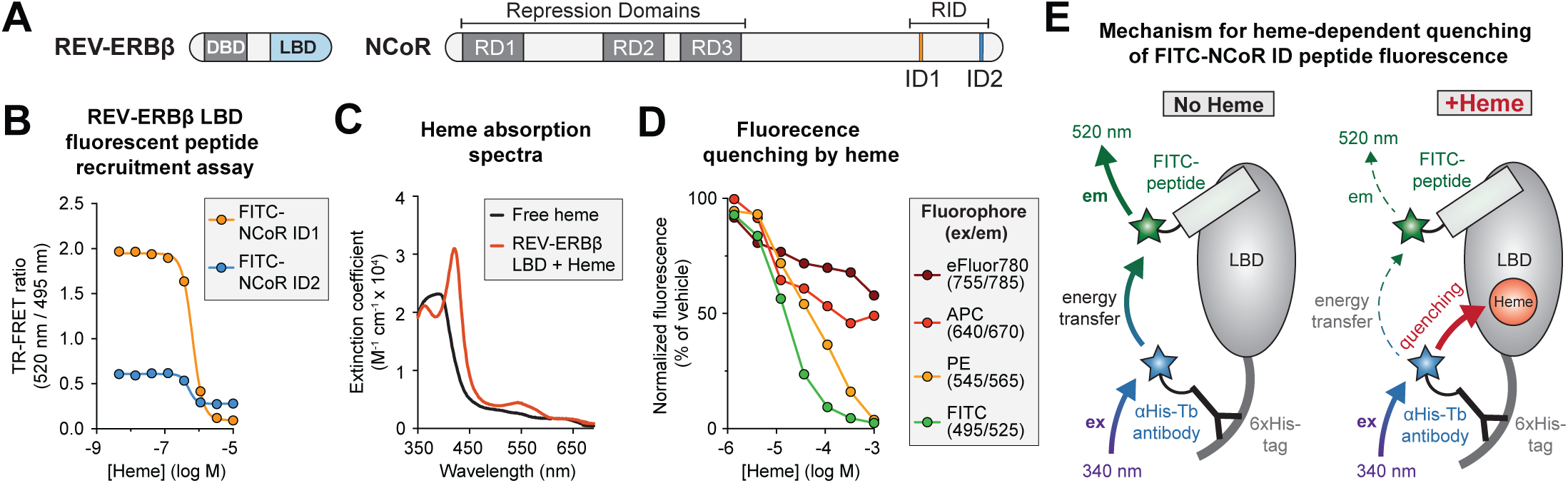
Heme-dependent fluorescence quenching in REV-ERB biochemical assays. (**A**) The conserved NR domain architecture includes a DNA binding domain (DBD) and ligand binding domain (LBD). NCoR is comprised of Repression Domains (RD) that mediate interactions with other transcriptional machinery and a nuclear receptor interaction domain (RID), which encompasses the interaction domain (ID) motifs ID1 and ID2. (**B**) TR-FRET assay to assess the interaction of FITC-conjugated NCoR ID peptides with REV-ERBβ LBD in response to a heme dose-response; a larger TR-FRET ratio indicates higher affinity binding and vice a versa (n=3; mean ± s.d.). (**C**) UV-visible absorption spectra of free heme and heme-bound REV-ERBβ LBD. (**D**) Dose-dependent heme quenching of fluorophores; excitation (ex) and emission (em) wavelengths used are indicated in the legend (n=2; mean ± s.d.). (**E**) Schematic indicating how heme could quench fluorescence in a TR-FRET assay to cause an artificially lower TR-FRET ratio.

### Heme and NCoR peptides cobind to the REV-ERBβ LBD

To test how heme affects REV-ERBβ LBD affinity for ID peptides using a fluorescence-independent method, we performed isothermal titration calorimetry (ITC), which provides information about the affinity and the thermodynamic parameters of binding. Heme decreased the REV-ERBβ LBD affinity for the NCoR ID1 peptide (0.5 μM to 2 μM) but increased the affinity NCoR ID2 peptide (16 μM to 2 μM) (**Fig. 3A and Table S1**). Heme also remodeled the thermodynamic profile of NCoR ID peptide binding to REV-ERBβ LBD, changing the interaction from enthalpically-driven and exothermic for apo-REV-ERBβ LBD to entropically-driven and endothermic for heme-bound REV-ERBβ LBD at 25°C (**Fig. 3B and Table S1**). These data show that heme-bound REV-ERBβ LBD displays equal affinity for NCoR ID1 and ID2 peptides and provide the first biochemical evidence that heme can directly increase REV-ERB affinity for NCoR.

**Fig. 3.**
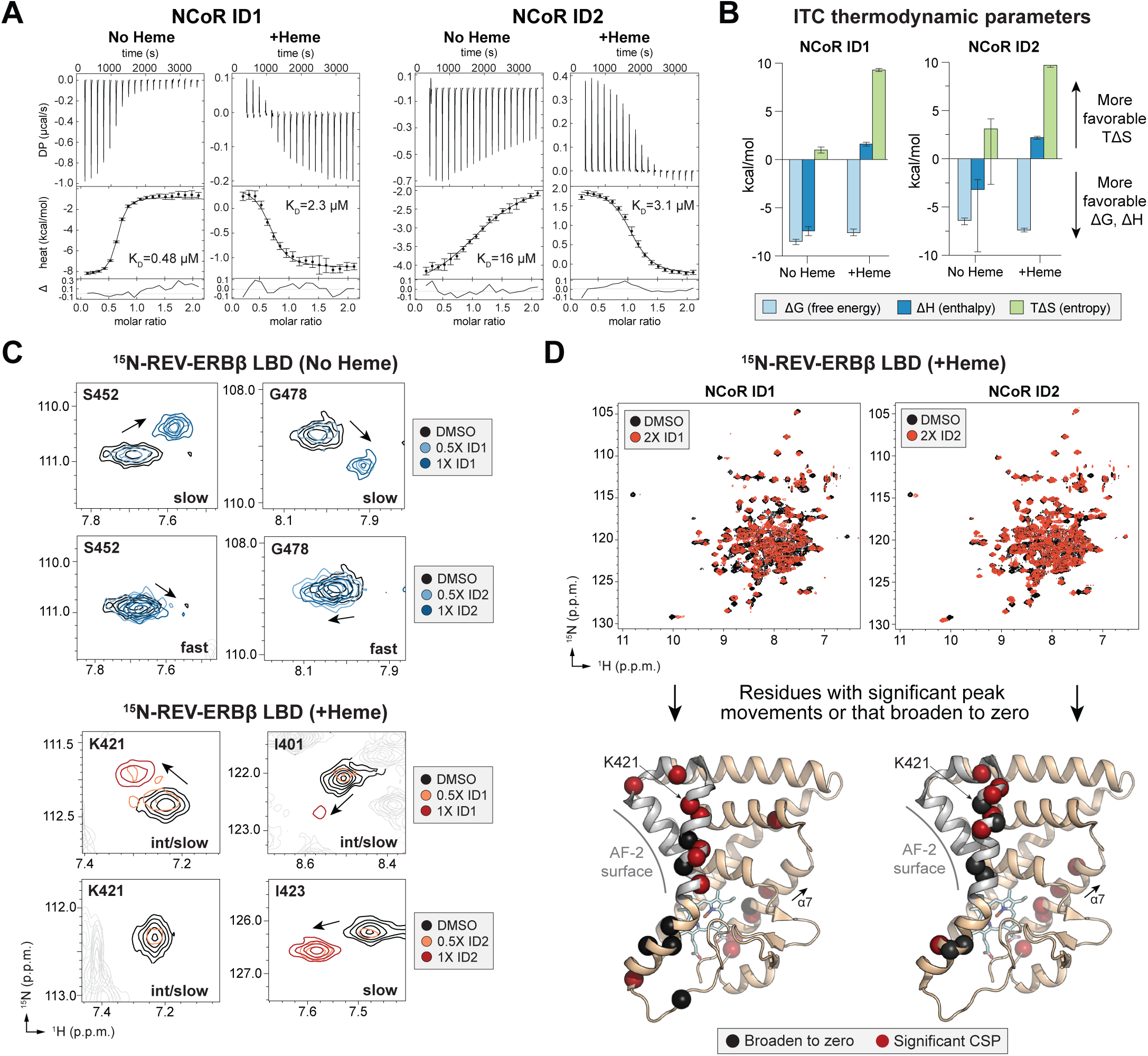
Heme can directly promote NCoR binding to REV-ERBβ LBD. (**A**) Representative ITC thermograms and fitted curves of REV-ERBβ LBD titrated with NCoR ID1 or ID2 peptide in the presence or absence of heme with calculated binding affinities inset in each plot (error bars, uncertainty in each injection calculated by the NITPIC algorithm). (**B**) Thermodynamic parameters calculated from global fitting analysis of replicate (*N* ≥ *2*) ITC experiments (mean with 95% confidence intervals calculated for each parameter as reported in **Table S1**). (**C**) Representative peaks from 2D [^1^H,^15^N]-TROSY-HSQC spectra of ^15^N-labeled REV-ERBβ LBD with or without heme titrated with NCoR ID1 or ID2 peptide. (**D**) 2D [^1^H,^15^N]-TROSY-HSQC spectral overlays of heme-bound REV-ERBβ LBD in the presence of vehicle (0.8% DMSO; black peaks) or 2 molar equivalents of ID1 or ID2 peptide (red peaks). Residues with the largest CSP changes or line broadening were mapped as red or black spheres, respectively, onto the crystal structure of heme-bound REV-ERBβ LBD (PDB 3CQV).

To support our ITC results, we performed solution NMR spectroscopy—a structural technique that provides information about the dynamics and structure (via changes to the chemical environment of individual atoms) in a protein at atomic resolution. Titration of NCoR ID peptide into ^15^N-labeled REV-ERBβ LBD can 1) validate that the interaction occurs in the presence of heme, 2) qualitatively reflect the strength of the interaction relative to apo (ligand-free)-REV-ERBβ LBD, and 3) highlight the residues most likely involved in binding (*16*). We previously obtained backbone NMR chemical shift assignments for REV-ERBβ LBD (*17*). Here, we collected 3D NMR data to obtain backbone NMR chemical shift assignments for heme-bound REV-ERBβ LBD. We confidently assigned 90 of 198 residues (**Fig. S1**); we were unable to assign the remainder likely due to intermediate exchange timescale dynamics in the heme-bound form and/or paramagnetic relaxation effects from the heme iron (*18*), both of which cause NMR peaks to broaden significantly.

Titration of NCoR ID1 or ID2 peptide into ^15^N-labeled REV-ERBβ LBD in the presence or absence of heme revealed localized NMR chemical shift changes (i.e., peak movements) in 2D [^1^H,^15^N]-TROSY-HSQC NMR spectra (**Fig. S2**), confirming the NCoR ID peptides bind to heme-bound REV-ERBβ LBD. Consistent with the higher ID1 affinity measured by ITC for apo-REV-ERBβ LBD (0.5 μM) compared to the heme-bound form (2 μM), ID1 peptide titration in the absence of heme displayed peak movements in slow exchange on the NMR timescale, which is characteristic of strong binding (**Fig. 3C and Fig. S2A**). For ID1 and ID2 peptides in the presence of heme, we observed a mixture of peak movements in slow and intermediate exchange on the NMR timescale (**Fig. 3C and Fig. S2, C and D**), which is typical of a moderate affinity binding event and agrees with the lower ID1 and ID2 peptide affinities in the presence of heme (2-3 μM). In the absence of heme, the ID2 showed only fast-to-intermediate exchange binding events (**Fig. 3C and Fig. S2B**), which is typical of moderate to weak binding and agrees with ITC data showing ID2 binding to apo-REV-ERBβ LBD was the weakest interaction (16 mM) (**Fig. 3A and Table S1**).

In other NRs, the AF-2 coregulator interaction surface is formed by the physical interaction of helix 12 with a surface formed by helices 3-5 within the LBD. REV-ERBβ lacks helix 12 but contains the surface formed by helices 3-5 including a conserved lysine residue (K421) on helix 3 important for corepressor interaction (*19*). To determine whether the NCoR ID motif peptides bind at the REV-ERBβ AF-2 surface in the presence of heme, we quantitatively analyzed chemical shift perturbation (CSPs) and NMR peak line broadening caused by peptide binding in the 2D [^1^H,^15^N]-TROSY-HSQC NMR titration spectra. In the presence of heme, AF-2 surface residues including K421 showed the largest CSP changes and line broadening, indicating the NCoR ID peptides bind at this surface (**Fig. 3D and Fig. S3**). We also observed binding effects on helix 7, which forms part of the ligand-binding pocket, which we attributed to peptide binding at the AF-2 surface allosterically inducing a shift in the conformation of heme and its propionate group adjacent to helix 7 (**Fig. S4**).

Collectively, our absorption/fluorescence, ITC, and NMR data support the notion that the heme-dependent complete displacement of NCoR ID peptides observed in fluorescence-based recruitment assays is an artifact of heme-dependent signal quenching. Our data further show that heme can directly enhance REV-ERBβ LBD affinity for the NCoR ID2 motif, resulting in equal affinity for both ID1 and ID2 motifs, and indicate the NCoR ID peptides bind at the AF-2 surface of heme-bound REV-ERBβ LBD.

### Crystal structures of REV-ERBβ LBD cobound to heme and NcoR peptides

In addition to fluorescence-based biochemical assays showing heme displaces NCoR ID peptides from the REV-ERB LBD, previously published structural data suggested heme and NCoR cobinding would be incompatible (*10, 11*). Overlay of crystal structures of apo-REV-ERBα LBD bound to NCoR ID1 peptide and heme-bound REV-ERBβ LBD without NCoR peptide indicated a kink in helix 3 of the heme-bound conformation would clash with the antiparallel β-sheet formed between NCoR ID1 peptide and a β-strand extension off helix 11 in the apo-conformation (**Fig. 1**). There is no published structure of apo-REV-ERBβ LBD bound to NCoR ID1 peptide, and we were unsuccessful in producing crystals of this complex; however, four lines of evidence suggest the NCoR ID1 β-sheet is present in REV-ERBβ. First, REV-ERBα and REV-ERBβ have high sequence conservation in β-strand extension off helix 11 (Blast, 100% positives). Second, the crystal structure of RARα LBD bound to NCoR ID1—the only other published NCoR ID1-bound NR LBD structure—also revealed an antiparallel β-sheet interaction with a C-terminal extension off helix 11 in RARα (*20*), indicating this may be a conserved NR/NCoR ID1 binding mode. Third, the apo-REV-ERBβ LBD affinity for NCoR ID1 (0.48 μM; 95% CI 0.31 – 0.73 μM) we determined by ITC is similar to apo-REV-ERBα LBD affinity for ID1 (0.4 ± 0.02 μM) also determined by ITC (*21*), indicating the contacts necessary for high affinity ID1 binding are conserved. Fourth, the NMR peaks for REV-ERBβ G478 and H575, residues structurally proximal to putative β-strand extension off helix 11, show a larger CSP when NCoR ID1 is titrated into apo-REV-ERBβ LBD compared to NCoR ID2 (**Fig. 3C and Fig. S2, A and B**), which is not predicted to form a β-strand (*10*).

Based on our ITC and NMR data that show heme and NCoR ID peptides can cobind the REV-ERBβ LBD, we hypothesized that structural changes in either the NCoR ID peptide—relative to conformation captured in the apo-REV-ERBα crystal structure—or in the conformation of heme-bound REV-ERBβ LBD without NCoR peptide would permit cobinding. We screened for crystals of the tertiary complexes and solved two structures of REV-ERBβ LBD cobound to heme and either NCoR ID1 peptide (2.6 Å resolution) or ID2 peptide (2.0 Å resolution) (**Fig. 4 and Table S2**). In both structures, REV-ERBβ LBD crystallized with two molecules in the asymmetric unit with an NCoR ID peptide and heme molecule cobound to each LBD. The overall RMSD of the REV-ERBβ LBD complex in the asymmetric unit, including LBD, heme, and NCoR ID peptide, was low—1.13 and 1.31 Å, respectively, for the ID1- and ID2-bound structures—suggesting minimal conformational heterogeneity. Consistent with our NMR data, the NCoR ID peptides were bound to the AF-2 surface (**Fig. 4**) and formed electrostatic interactions with the conserved charge clamp residue K421 on helix 3, confirming the heme-dependent REV-ERBβ corepressor interaction is similar to other NRs (*19*). These structures allowed us to explore structural changes in the heme and NCoR ID cobound state vs. the ID peptide- or heme-only bound structures.

**Fig. 4.**
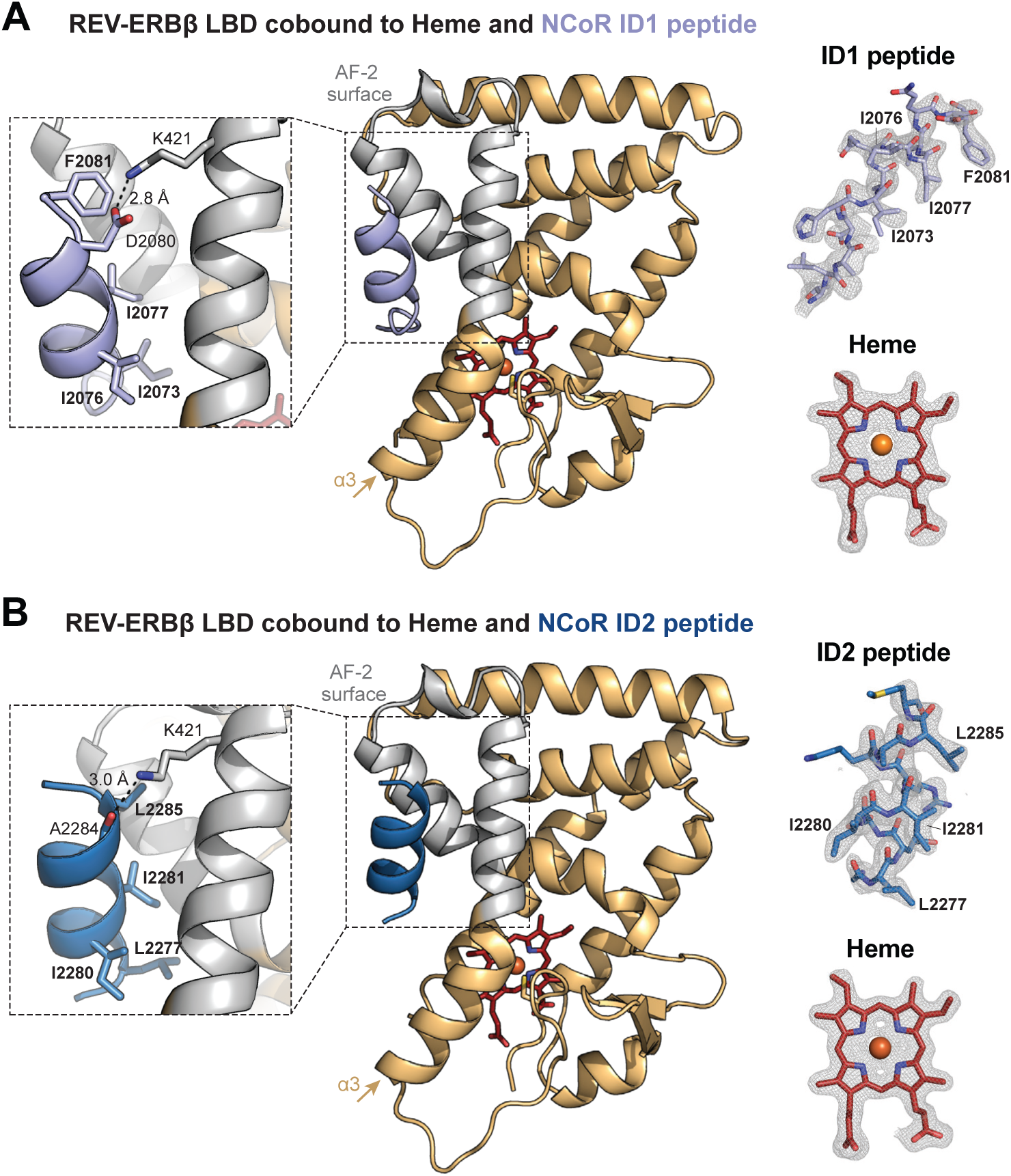
Crystal structures of REV-ERBβ LBD cobound to heme and NCoR ID peptides. (**A**) Structure of REV-ERBβ LBD (light orange cartoon with the AF-2 surface in gray) cobound to heme (red sticks) and NCoR ID1 peptide (light purple cartoon) (PDB 6WMQ). (**B**) Structure of REV-ERBβ LBD (light orange cartoon with the AF-2 surface in gray) cobound to heme (red sticks) and NCoR ID1 peptide (dark blue cartoon) (PDB 6WMS). Insets to the left of the structures highlight the ID peptide CoRNR box motif residues (in bold) and conserved charge clamp interaction with K421. Omit maps (2F_o_-F_c_, contoured at 1σ) for the ID peptides and heme are shown to the right of the structures with CoRNR box motif residues indicated.

Relative to the previously published hemebound REV-ERBβ LBD structure (*11*), the overall LBD conformations were largely unchanged by peptide binding with an average RMSD of 1.00 Å for the ID1 structure and 1.25 Å for the ID2 structure. Since the heme-bound LBD conformation was unchanged, the NCoR ID1 peptide binding mode was necessarily altered to accommodate cobinding: the antiparallel β-sheet between the N-terminus of NCoR ID1 and the β-strand extension off helix 11 did not form in our heme-bound REV-ERBβ LBD structures (**Fig. S5A**). Although the antiparallel β-sheet was lost, the interaction with the C-terminal α-helical region of the NCoR ID1 and ID2 peptides containing the I/LxxI/LI/LxxxI/L/F CoRepressor Nuclear Receptor (CoRNR) box motif critical for corepressor binding (*10*) was preserved. These data indicate that heme stabilizes a REV-ERBβ LBD conformation that alters the NCoR ID1 antiparallel β-sheet binding mode but preserves the α-helical CoRNR box motif interaction.

These structural features in the heme and NCoR ID cobound crystal structures are supported by our ITC data. Heme decreased REV-ERBβ LBD affinity for ID1, which is consistent with heme-dependent inhibition of antiparallel β-sheet formation and elimination of contacts that contribute to higher affinity binding. This conclusion is supported by a previous study showing truncation of β-sheet-forming NCoR ID1 residues reduced apo-REV-ERBα affinity from 0.4 to 5 μM: the magnitude of this effect is nearly identical to the heme-dependent reduction in ID1 affinity (from 0.5 to 2 μM) we observed (*21*). Despite loss of the antiparallel β-sheet interaction, the essential AF-2 surface interactions with the α-helical CoRNR box motif are preserved, causing heme to weaken—but not eliminate— ID1 binding. This theory aligns well with the similar affinities for NCoR ID1 and ID2 to heme-bound REV-ERBβ LBD (both ∼2-3 μM) (**Table S1**) and the overlapping α-helical binding modes observed for NCoR ID1 and ID2 peptides (the overall RMSD between the structures was 1.66 Å) (**Fig. S5B**). Thus, our crystal structures explain the heme-dependent decrease in ID1 affinity observed by ITC.

Our crystal structures also offer insight into the heme-dependent increase in ID2 affinity observed by ITC. Likely due to its weaker affinity, we were unsuccessful in generating crystals of NCoR ID2 bound to apo-REV-ERBβ LBD. However, published structures of other NR LBDs bound to NCoR ID2 peptides showed that ID2 does not form the antiparallel β-sheet interaction with NR LBDs observed for ID1 (*19*). This could explain why NCoR ID2 had the weakest affinity to apo-REV-ERBβ LBD: if the antiparallel β-sheet interaction is important for NCoR ID binding to apo-REV-ERB LBD, but NCoR ID2 does not contain a β-strand, the ID2 would be incapable of strong binding in the absence of heme. On the other hand, heme appears to be necessary to strengthen binding of the α-helical CoRNR box within the ID motifs, overcoming the need for β-sheet formation with the REV-ERBβ LBD. We speculate that the heme-dependent increase in NCoR ID2 affinity is likely due to heme, which is a highly hydrophobic molecule, enhancing the overall hydrophobicity of the pocket and nearby AF-2 surface, which concomitantly drive higher affinity NCoR CoRNR box motif binding. A central role for the hydrophobic effect is supported by the large gain in entropy observed for heme-bound REV-ERBβ LBD in our ITC assays for both ID1 and ID2; we attribute the favorable entropic forces to desolvation of the hydrophobic AF-2 surface upon ID binding (*22*). Together, these data support a model in which heme primes the REV-ERBβ LBD for more favorable CoRNR box motif binding by enhancing AF-2 hydrophobic interactions.

### Heme enhances REV-ERBβ LBD affinity for the NCoR RID

Our ITC data showed that heme alters REV-ERBβ affinity for ID peptides such that ID1 and ID2 affinity in the presence of heme are approximately equal. We hypothesized that this might be advantageous in the context of the disordered NCoR RID containing both ID motifs (*23*) (**Fig. 2A**). Strikingly, ITC experiments showed that heme significantly increased REV-ERBβ LBD affinity for the NCoR RID by about 4-fold (350 nM to 88 nM) (**Fig. 5A and Table S3**). Consistent with our ITC data using the individual NCoR ID peptides, heme altered the thermodynamic profile of RID binding from enthalpically-driven and exothermic to entropically-driven and endothermic at 25°C (**Fig. 5B and Table S3**). This consistency in the thermodynamic profiles for the RID and individual ID peptides suggests that the individual ID motifs within the RID are most likely mediating binding. Additionally, the heme-dependent increase in affinity and altered thermodynamic profile was accompanied by a change in the REV-ERBβ:NCoR RID interaction stoichiometry from 1:1 (no heme) to 2:1 (+heme) (**Fig. 5A and Table S3**).

**Fig. 5.**
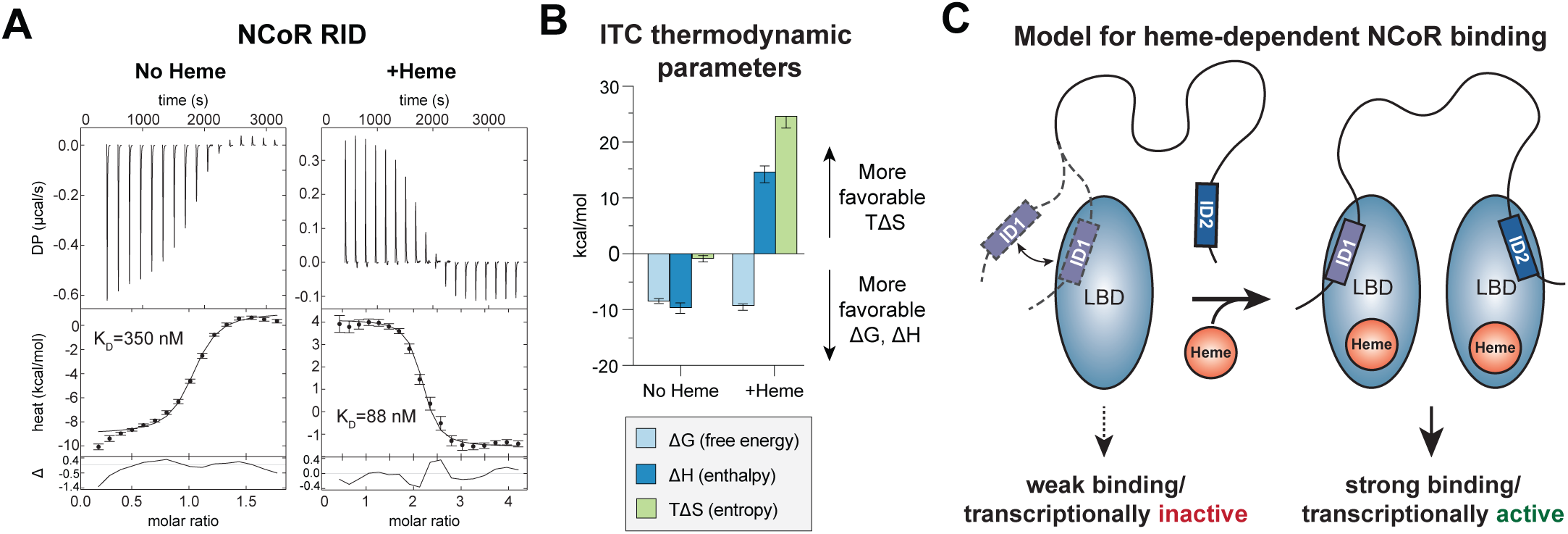
Heme increases REV-ERBβ LBD affinity for an NCoR RID construct comprised of both ID motifs. (**A**) Representative ITC thermograms and fitted curves of REV-ERBβ LBD and NCoR RID in the presence or absence of heme with calculated binding affinities highlighted in red (error bars, uncertainty in each injection calculated by the NITPIC algorithm). (**B**) Thermodynamic parameters calculated from global fitting analysis of replicate (*N ≥ 2*) experiments in (**A**) (mean ± 95% confidence intervals calculated for each parameter as reported in **Table S3**). (**C**) Schematic indicating a model for heme-dependent REV-ERB interaction with NCoR RID. The affinity of the ID1 motif alone is insufficient to drive binding of NCoR RID to apo-REV-ERB. However, in the presence of heme, cooperative binding of both NCoR ID motifs promotes high affinity binding and permits active repression of REV-ERB-dependent transcription.

Taken together, the similar NCoR RID and ID1 peptide affinities (350 and 480 nM, respectively) indicate that the ID1 motif drives NCoR RID to binding to apo-REV-ERBβ LBD, Moreover, the molecular basis for the heme-dependent increase in NCoR affinity arises through cooperative binding of both ID motifs within the NCoR RID to one REV-ERBβ LBD (**Fig. 4C**). Intriguingly, this 2:1 stoichiometry agrees with a previous study showing that REV-ERBα could only interact with full-length NCoR on DNA response elements containing two REV-ERB binding sites (*24*), suggesting REV-ERB interaction with NCoR could be governed by heme’s ability to remodel the stoichiometric properties of binding. Importantly, our data support cellular studies showing heme is required for REV-ERBβ to interact with NCoR (*7-9*): the heme-dependent increase in affinity could dictate whether NCoR binds REV-ERBβ in a cellular environment where protein concentrations are limiting.

## DISCUSSION

The closely related REV-ERB nuclear receptors, REV-ERBα and REVERBβ, are transcriptional repressors whose natural ligand is the iron-centered porphyrin heme. Heme’s role in regulating REV-ERB activity has been unclear due to contradictory cell-based vs. structural evidence (*7-11*). Here, we resolve this conflict by providing evidence that heme directly increases REV-ERBβ affinity for the transcriptional corepressor protein NCoR. We present fluorescence-independent binding data and two crystal structures that reveal the structural basis for heme and NCoR cobinding to REV-ERBβ. Mechanistically, our results indicate that heme enhances hydrophobic interactions to facilitate cooperative binding of the NCoR ID motifs. By revealing the biochemical and structural basis for previously reported heme-dependent functional observations in cellular studies, our data support a cohesive model in which the REV-ERBs are transcriptionally inactive without heme, which is required to bind NCoR and actively repress transcription. Collectively, our results should facilitate renewed efforts toward understanding the structure and function of ligand-regulated REV-ERB activity in health and disease.

### Structural features mediating NCoR binding to REV-ERB LBD

Of the two published crystal structures of NR LBDs bound to NCoR ID1 peptide, both show that ID1 induces an α-helix to β-strand secondary structure switch in the C-terminal extension off helix 11 resulting in an antiparallel β-sheet interaction with a β-strand region of NCoR ID1 (*10, 20*). Our structure of REV-ERBβ LBD cobound to heme and NCoR ID1 did not form this antiparallel β-sheet interaction, demonstrating that it is dispensable for ID1 binding in the presence of heme. While elimination of β-sheet formation by truncation or mutation of the NCoR ID1 peptide or β-strand-forming residues in RARα drastically reduced RARα affinity for NCoR ID1 (∼100-fold) and abolished NCoR-RARα interaction in cells (*20*), loss of the β-sheet only partially reduced REV-ERBα LBD binding to NCoR ID1 (2.5- to 10-fold) (*10, 21*). It was speculated that formation of the antiparallel β-sheet between RARα LBD and NCoR ID1 promotes corepressor binding by displacing the AF-2 helix 12, which competes with corepressors for docking at the AF-2 surface (*20*); however, this function is nonessential in REV-ERBs, which lack a helix 12, explaining why loss of the antiparallel β-sheet interaction is not as detrimental to ID1 interaction with the REV-ERBs. Nonetheless, evidence indicates that both apo-REV-ERBα and apo-REV-ERBβ can form the antiparallel β-sheet with NCoR ID1, calling into question the functional role of this interaction. Given that heme is required for NCoR binding in cells (*7-9*) and that heme disrupts the antiparallel β-sheet, one possibility is that the REV-ERB/ID1 β-sheet interaction is an evolutionary artifact and therefore not functionally relevant; i.e., the antiparallel β-sheet interaction only occurs under supraphysiological protein concentrations and/or for apo-REV-ERBs in the context of the isolated ID1 peptide. Alternatively, the antiparallel β-sheet interaction may be important for forming a REV-ERB-NCoR encounter complex that facilitates access of the α-helical NCoR ID1 CoRNR box motif to the REV-ERB AF-2 surface prior to heme binding and concomitant stabilization of CoRNR box interactions (**Fig. 5C**). Cellular protein-protein interaction assays, such as mammalian two-hybrid or co-immunoprecipitation, in which the NCoR ID1 β-strand residues are mutated are needed to address these possibilities.

### Stoichiometric principles of REV-ERB-dependent transcriptional repression

The REV-ERBs are unique among NRs because they lack the C-terminal AF-2 helix important for binding transcriptional coactivator proteins; thus, the REV-ERBs are thought to exclusively bind transcriptional corepressor proteins and repress transcription (2). This apparent capacity for constitutive repression calls into question how REV-ERB activity is regulated, since unrestrained transcriptional repression would be unfavorable in most cells. One possibility that has been explored is that REV-ERB DNA binding stoichiometry dictates REV-ERB activity. The REV-ERBs share a monomeric DNA response element with the ROR NRs known as an ROR response element, or RORE (*25*). Since the RORs generally activate transcription, REV-ERB binding to ROREs prevents RORs from binding and recruiting transcriptionally activating complexes that modify chromatin (i.e., increase acetylation) to enhance access of RNA polymerase and its associated machinery (*26*). This process in which REV-ERBs sterically block RORs from activating their shared target genes is known as passive repression. However, the REV-ERBs are also capable of active repression. Active repression occurs when the REV-ERBs recruit transcriptionally repressive complexes that modify chromatin (i.e, increase deacetylation) to prevent access of RNA polymerase and its associated machinery (*26*). Previous studies have shown that the REV-ERBs can only actively repress transcription at sites containing two ROREs, referred to as REV-ERB response elements or REV-REs (*5, 27*). The molecular basis for this repressive mechanism is a result of REV-ERB interaction with the NCoR corepressor on REV-RE sites but not monomeric RORE sites (*24*). These findings demonstrate that stoichiometric principles govern REV-ERB-dependent effects on target gene expression.

Although DNA-dependent stoichiometry may be one mechanism influencing REV-ERB activity, the REV-ERBs are in the NR family that evolved to be ligand-regulated— in principle, the natural REV-ERB ligand heme should be important for regulating REV-ERB function. Cellular studies indicated that heme is required for REV-ERB interaction with NCoR and active transcriptional repression (*7-9*). Taken together with the stoichiometric findings discussed above, these data indicate that REV-ERBs require both DNA binding to a two site REV-RE and heme binding to the LBD to recruit NCoR and actively repress transcription. This raises a compelling question as to whether there is a connection between heme binding and DNA-dependent REV-ERB:NCoR stoichiometry on REV-RE sites but not monomeric RORE sites. Our ITC results using the NCoR RID showed that heme binding changes the REV-ERBβ:NCoR stoichiometry from 1:1 to 2:1, suggesting heme could be important for increasing NCoR affinity specifically at REV-RE sites. A previous study characterized the interaction of full-length REV-ERBβ with the NCoR RID on REV-RE response elements by EMSA using FAM-labeled DNA (*9*). REV-ERBβ interacted with RID on these multi-site REV-REs in a concentration-dependent manner, consistent with observations for REV-ERBα (*24*) and the relatively high affinity of apo-REV-ERBβ for NCoR RID (**Fig. 5A**). However, although stoichiometric heme permitted NCoR binding on DNA, increasing heme concentrations appeared to dissociate the interaction of NCoR RID with both REV-ERBβ (a heme-binding protein) and THRβ (a NR that does not bind heme) on FAM-labeled DNA (*9*). This was attributed to potential Fenton chemistry-related effects, whereby high concentrations of free heme may interfere with DNA binding of both NRs either through direct interaction of heme with DNA or by heme inducing DNA degradation; however, our data suggest the effect may be due to heme quenching the fluorescent FAM-labeled DNA signal. Future studies are warranted to further explore the interplay between DNA binding and heme-dependent NCoR interaction using ITC or a fluorescence-independent EMSA assay (e.g., using radiolabeled DNA). Our ITC data suggest heme should promote a higher affinity interaction, which importantly could distinguish whether binding occurs in cells where REV-ERBβ and NCoR protein concentrations are concievably much lower. Indeed, additional evidence from that study supports this idea, since REV-ERBβ mutants incapable of binding heme interacted with NCoR in the EMSA assay, but did not coimmunoprecipitate with NCoR in cellular extracts (*9*). To expand upon these biochemical studies, we further propose that structural methods such as SAXS would help address whether heme promotes NCoR binding at multiple REV-ERB LBD molecules on DNA, as suggested by our ITC data. Finally, an important question that remains to be addressed is how heme regulates REV-ERB function at monomeric RORE sites, where heme should be capable of binding, but NCoR should not be recruited based on current evidence.

### Conserved heme-dependent mechanisms for REV-ERBα and REV-ERBβ

One caveat to our work is that it focuses on REV-ERBβ, so the relevance of our conclusions to REV-ERBα are not guaranteed. REV-ERBα and REV-ERBβ are thought to be functionally redundant, and cell-based studies have reported a number of similarities including heme-dependent NCoR binding and overlapping target genes (*7-9, 28*). Indeed, the DNA-binding domain (DBD) sequences of both REV-ERBs are 95% identical (Blast, 100% positives) and the core LBD sequences (helix 3 – C-terminus) are 76% identical (Blast, 88% positives). It is also compelling that NCoR ID peptide affinities we determined for apo-REV-ERBβ LBD (**Table S1**) are in excellent agreement with apo-REV-ERBα (*21*). Despite the similarities in the REV-ERB LBD sequences, recombinant REV-ERBα LBD displays poor solubility that is prohibitive to structural studies—unlike REV-ERBβ LBD, which is very soluble. A REV-ERBα LBD deletion construct (residues 281-614Δ324-422) was previously generated with sufficient solubility to solve the crystal structure of apo-REV-ERBα LBD bound to NCoR ID1 (*10*). However, we found this construct, as well as the full REV-ERBα LBD (residues 281-614), precipitated in the presence of heme, regardless of buffer, which could explain why that previous study was unable crystallize the REV-ERBα LBD construct cobound to heme and NCoR (*10*). Heme-dependent REV-ERBα LBD precipitation and/or fluorescence quenching effects could also explain why a previous study reported reduced REV-ERBα LBD affinity for an NCoR RID construct in the presence of heme using fluorescence polarization assays (*21*). Ultimately, although current evidence strongly suggests REV-ERBα and REV-ERBβ function similarly, future studies are warranted to confirm that heme-dependent structural mechanisms we report here for REV-ERBβ also apply to REV-ERBα.

### Biological significance of heme as an endogenous REV-ERB ligand

Heme has a diverse repertoire of essential functions. Heme’s role as an enzyme cofactor in delivery of diatomic gases, lipid/xenobiotic oxidation, electron transfer in oxidative metabolism, etc. are well-documented. However, recent studies have established that free, or labile, heme functions as a signaling molecule (*29*). Several intracellular heme transporters have been reported including PGRMC2, which was shown to be essential for delivering heme synthesized in mitrochondria to the nucleus, where it binds heme-responsive transcription factors including REV-ERBα (*30*). Despite identification of this heme transport pathway, several important questions remain. In this study, we have addressed one of these questions by providing structural data necessary support a model in which heme directly enhances corepressor recruitment to REV-ERBβ to repress transcription. Our results should facilitate renewed investigations into the remaining questions concerning the biological significance of heme as a REV-ERB-dependent signaling molecule. For example, although heme is known to signal through the REV-ERBs, the information conveyed through heme is not well understood. One possibility is that signaling pathways that upregulate heme biosynthesis concomitantly activate REV-ERB-dependent target genes as part of a feedback mechanism. Future studies are needed to explore the pathways that modulate heme biosynthesis (i.e., by regulating the expression of heme enzymes or mitochondrial biogenesis) and their potential effects on REV-ERB activity. The results of such studies should provide exciting new insight into heme-dependent REV-ERB biology.

## MATERIALS AND METHODS

### Plasmids and reagents

The human REV-ERBβ ligand binding domain (LBD; residues 381-579) was previously cloned into the pET46 vector (*17*). The mouse NCoR Receptor Interaction Domain (RID; residues 1942-2208 in the X50 splice variant), which has 87% sequence identity to human NCoR RID (100% sequence identity in the ID motifs), was previously cloned into the pET32 vector (*23*). Heme (Sigma #51280) was prepared either as a 1 mM stock in DMSO stored at 4°C or 50 mM stock in 0.2M NaOH (solutions prepared in 0.2M NaOH were always made fresh immediately prior to the experiment), where indicated. NCoR ID1 peptide (RTHRLITLADHICQIITQD-FARN) and NCoR ID2 peptide (DPASNLGLEDIIRKALMGSFDDK) were purchased with >95% from Lifetein with N-terminal amidation and C-terminal acetylation for stability and prepared as 50 mM stocks in DMSO stored at −80°C.

### REV-ERBβ LBD expression and purification

Human REV-ERBβ LBD was expressed in BL21/DE3 *Escherichia coli* cells with an N-terminal hexahistidine (6xHis) tag separated by a 3C protease cleavage site. Expression was performed using autoinduction media: cells were grown at 37°C for 5 hours, 30°C for 1 hour, and 18°C for 16 hours before harvesting by centrifugation. Pellets were resuspended in potassium phosphate lysis buffer (40 mM potassium phosphate, pH 7.4, 500 mM KCl, 15 mM imidazole, 1 mM DTT) supplemented with leupeptin, pepstatin A, lysozyme, and DNase I and sonicated on ice. Lysed cells were centrifuged 20,000 xg for 30 minutes at 4°C and soluble lysate was filtered prior to IMAC purification using 2×5mL His-Trap columns (GE Health-care) affixed to an Akta Pure. A 5 mL aliquot of 6xHis-tagged protein was purified by size exclusion chromatography using a Superdex 75 column equilibrated in TR-FRET assay buffer (20 mM potassium phosphate, pH 7.4, 50 mM KCl, 0.5 mM EDTA); purified aliquots were stored at −80°C for TR-FRET assays. To cleave the 6xHistag from the remaining protein, it was incubated overnight with 3C protease (generated in-house) via dialysis; then, protein was reloaded onto the His-Trap columns and the flow-through collected. Finally, aggregates were removed by size exclusion chromatography using a Superdex 75 column and HEPES gel filtration buffer (20 mM HEPES, pH 7.4, 50 mM KCl, 0.5 mM EDTA). >95% purity was confirmed by SDS-PAGE. Protein was aliquoted and stored at −80°C.

### Crystallization, data collection, and structure determination

REV-ERBβ LBD was incubated with 2 molar equivalents of heme (prepared fresh at 50 mM in 0.2M NaOH) overnight at 4°C before addition of 5 molar equivalents of either NCoR ID1 peptide or NCoR ID2 peptide. After overnight incubation at 4°C with peptide, REV-ERBβ LBD cobound to heme and peptide was buffer exchanged into HEPES gel filtration buffer to remove unbound and concentrated to 15 mg/mL (ID2-bound) or 5 mg/ mL (ID1-bound). Buffer exchanged proteins were screened for conditions that produced crystals using the NR LBD (Molecular Dimensions), Structure (Molecular Dimensions), INDEX (Hampton Research), and PEG Ion (Hampton Research) kits and sitting drop method (1 μL reservoir solution added to 1 μL drop of protein solution) at 22°C. REV-ERBβ LBD crystals cobound to heme and NCoR ID2 grew in 0.2 M Mg formate dihydrate, 20% w/v PEG 3350 (PEG Ion) and REV-ERBβ LBD crystals cobound to heme and NCoR ID1 grew in 2.0 M ammonium sulfate, 0.1 M Na HEPES, pH 7.5, 2% PEG 400 (Structure). Crystals in their respective condition were supplemented with 10% glycerol and flash cooled in liquid nitrogen. X-ray diffraction data were collected at the ALS synchrotron (Lawrence Berkeley National Laboratory) in 180 images with a 1-degree rotation per image. Data were processed, indexed, and scaled using Mosflm and Scala in CCP4 (*31, 32*). Molecular replacement was performed using the program Phaser (*33*) in the PHENIX software package (*34*) using the heme-bound REV-ERBβ LBD crystal structure as a search model (PDB 3CQV) (*11*). The structures were solved at 2.55 Å (for the ID1-bound structure) and 2.0 Å (for the ID2-bound structure). The NCoR ID1 and NCoR ID2 peptides were manually built into the unmodelled density using COOT (*35*). Subsequent iterations of automated refinement were performed using PHENIX with additional manual refinement in COOT.

### Structural alignments and RMSD calculations

Structures were aligned and RMSD values were calculated in PyMOL using the “align” command with cycles set to 0 (no outliers were removed). Waters were excluded from all alignments. For the alignment of the chains within the asymmetric units, LBD, heme and peptide were aligned together as a single object. For alignment of the heme and peptide cobound structures with the published heme-bound structure, heme and LBD were included, while peptide was excluded. Both chains in the heme and NCoR ID peptide cobound REV-ERBβ LBD structures were aligned individually to the heme-bound REV-ERBβ LBD (which crystallized as monomer) and the average of the two values was reported. For alignment of the heme-bound REV-ERBβ LBD cobound to NCoR ID1 and ID2—the LBD, heme, and ID peptide were aligned together as a single object; the respective chains A and B were aligned and the average RMSD of the two values was reported.

### TR-FRET

6xHis-tagged REV-ERBβ LBD (4 nM) was incubated with anti-His antibody conjugated to terbium (1 nM) (ThermoFisher #PV5863) and FITC-labeled NCoR ID1 or NCoR ID2 peptide (400 nM) in assay buffer (20 mM potassium phosphate, pH 7.4, 50 mM KCl, 0.5 mM EDTA). A 2:1 dose response of heme prepared as a 1 mM stock in DMSO was added to the protein mixture, mixed, and 20 μL/well was transferred to a black bottom 384-well plate (Greiner). After 2 hour incubation at 4°C, the plate was read using a Biotek Synergy plate reader (395 nm excitation; 495 nm terbium emission; and 520 nm FITC emission). The TR-FRET ratio was calculated as the fluorescence intensity at 520 nm divided by the intensity at 495 nm and plotted as a function of heme concentration in GraphPad Prism. The log-transformed data was fit to a sigmoidal dose response curve. The plot shown is representative of two independent experiments.

### Absorption spectra

Heme prepared as a 1 mM stock in DMSO was diluted to 20 μM in potassium phosphate buffer (20 mM potassium phosphate, pH 7.4, 50 mM KCl, 0.5 mM EDTA) and three wells of 200 μL/well were plated in a clear-bottom 96-well plate (Corning). 20 μM REV-ERBβ LBD was diluted in potassium phosphate buffer and mixed with 1.05 molar equivalents of heme and three wells of 200 μL/well were also plated in the clear-bottom 96-well plate. Absorbance from 350-700 nm was measured using a Biotek Synergy plate reader, values were averaged and subtracted from the spectrum of buffer only. The molar extinction coefficient at each wavelength was calculated given a path length of 1 cm and concentrations of 20 μM.

### Fluorescence quenching

A 1:2 dose response of heme (prepared as a 50 mM stock in 0.2M NaOH) was prepared in HEPES gel filtration buffer at 2X final concentration with a starting concentration of 2 mM. The heme dose response was mixed 1:1 with 2X-concentrated antibodies conjugated to fluorescent dyes (FITC; Biolegend #100510 and eFluor780; eBioscience #65-0865-14) or proteins (APC; Biolegend #103012 and PE; Biolegend #100206) prepared as 2 μg/ mL solutions in black-bottom 96-well plates (Greiner), with 100 μL total volume per well. Two wells per heme concentration were averaged as technical replicates. Fluorescence was analyzed using a Biotek Synergy plate reader and fluorescence intensity was plotted as a function of heme concentration in Graphpad Prism and normalized to the heme vehicle (DMSO) to facilitate comparison. For the FITC-conjugated antibody, the excitation wavelength was 425 nm and the emission wavelength was 525 nm. For the PE-conjugated antibody, the excitation wavelength was 545 nm and the emission wavelength was 565 nm. For the APC-conjugated antibody, excitation wavelength was 640 nm and the emission wavelength was 670 nm. For the eFluor780 dye, excitation wavelength was 755 nm and the emission wavelength was 785 nm. Data are representative of two individual experiments.

### Isothermal titration calorimetry

NCoR ID2 peptide was prepared at 1 mM in assay buffer (20 mM HEPES, pH 7.4, 50 mM KCl, 0.5 mM EDTA, 5 mM TCEP) and REV-ERBβ LBD was prepared at 100 μM with 2% DMSO to match the NCoR ID2 vehicle. NCoR ID2 in the syringe was titrated into REV-ERBβ LBD in the sample cell at 2 μL intervals at 25°C, with 60 second delay between injections for a total of 20 injections, or 2 molar equivalents of peptide (mixing speed of 1200 rpm) using a MicroCal iTC200. For experiments with heme, 1.05 molar equivalents of heme prepared in 0.2 M NaOH at 50 mM were added to the REV-ERBβ LBD and 0.2% 0.2M NaOH was added to the peptide to match the heme vehicle. NITPIC software (*36*) was used to calculate baselines, integrate curves, and prepare experimental data for unbiased global fitting analysis in SEDPHAT (*37*). *N = 2* experimental replicates without heme and *N = 3* experimental replicates with heme were fit to generate final binding affinity and thermodynamic parameter measurements. NCoR ID1 peptide was prepared at 50 mM in assay buffer and REV-ERBβ LBD was prepared at 500 μM with 0.1% DMSO to match the NCoR ID1 vehicle. REV-ERBβ LBD in the syringe was titrated into NCoR ID1 in the sample cell as described above for ID2. *N = 2* experimental replicates without heme and *N = 2* experimental replicates with heme were fit to generate final binding affinity and thermodynamic parameter measurements using NITPIC and SEDPHAT, as for ID2. NCoR RID was prepared in assay buffer at 50 μM and REV-ERBβ LBD was prepared at 500 μM. REV-ERBβ LBD in the syringe was titrated into the RID in the sample cell as described above for ID1 and ID2. *N = 2* experimental replicates without heme and *N = 2* experimental replicates with heme were fit to generate final binding affinity and thermodynamic parameter measurements using NITPIC and SEDPHAT, as for the ID peptides. Final figures were exported to GUSSI for publication-quality figure preparation (*38*). Control experiments were performed, including 500 μM REV-ERBβ LBD into assay buffer and 1 mM ID2 peptide into assay buffer, which revealed negligible heats of dilution, confirming that all observed signals were attributable to heats of binding.

### Generation of isotopically-labeled REV-ERBβ LBD

For generation of ^15^N-labeled REV-ERBβ LBD, protein was expressed in BL21/DE3 *Escherichia coli* cells using M9 minimal media supplemented with ^15^NH_4_Cl (Cambridge Isotope Laboratories) induced at an OD_600_ of 0.6 with 0.5 mM IPTG for 16 hours at 18°C. For generation of ^2^H, ^15^N,^13^C-labeled REV-ERBβ LBD (∼70% deuteration) for peak assignment experiments, cultures were first grown at 37°C in 1 L of LB media until an OD_600_ of 0.6 before the cells were pelleted and resuspended in 0.5 L M9 minimal media prepared in D_2_O supplemented with ^13^C-D-glucose and ^15^NH_4_Cl (Cambridge Isotope Laboratories). After 1 hour recovery at 37°C, expression was induced with 0.5 mM IPTG for 16 hour at 18°C before harvesting by centrifugation. The purifications were performed as described for unlabeled protein.

### NMR spectroscopy

To generate heme-bound protein, 1.25 molar equivalents of heme prepared in 0.2 M NaOH as a 50 mM stock were added to ^15^N-labeled REV-ERBβ LBD (200 μM) or ^2^H,^15^N,^13^C-labeled REV-ERBβ LBD (1 mM) in HEPES buffer (20 mM HEPES, pH 7.4, 50 mM NaCl, 0.5 mM EDTA). TROSY-based 3D HNCO, HNCA, HN(CA)CB, HN(COCA)CB, HN(-CO)CA, and HN(CA)CO experiments were collected at 298 K using the heme-bound ^2^H,^15^N,^13^C-REV-ERBβ LBD. 2D [^1^H,^15^N]-TROSY-HSQC data were collected for heme-bound ^15^N-labeled REV-ERBβ LBD at 298 K with addition of NCoR ID1 or ID2 peptides prepared as 50 mM stock in DMSO and analyzed in NMRViewJ. Data were collected on a Bruker 700 MHz NMR system equipped with a QCI cryoprobe, processed using NMRFx (*39*), and analyzed using NMRViewJ (*40*). Chemical shift perturbation (CSP) analysis was performed using 2D [^1^H,^15^N]-TROSY-HSQC heme-bound REV-ERBβ spectrum. Transfer of peak assignments from the vehicle (0.8% DMSO) to the 2X NCoR ID1- or ID2-bound spectra was performed using the minimal NMR chemical shift method (*16*). Peaks were identified to have broadened to zero if there was no confident peak in proximity to the vehicle peak. The average CSP and the standard deviation (SD) in the CSPs was calculated for NCoR ID1 or ID2 titrations, and the peaks that displayed CSPs in the presence of peptide greater than >1 SD above the average CSP were noted as significant.

### NCoR RID expression and purification

The mouse NCoR RID construct was transformed into BL21/DE3 *Escherchia coli* cells, which were grown in 2 L Terrific Broth (TB) media (24 g/L yeast extract, 12 g/L tryptone, 23.1 g/L KH_2_PO_4_, 125.4 g/L K_2_HPO_4_, and 4 mL/L glycerol) at 37°C until OD_600_ reached 0.7-0.8. Expression was induced with 0.75 mM IPTG for 2 hours at 37°C before cells were harvested by centrifugation. Pellets were washed 1X with PBS and stored −80°C until purification. Pellets were resuspended on ice in HEPES lysis buffer (25 mM HEPES, pH 7.4, 150 mM NaCl, 2 M urea, and 15 mM imidazole) supplemented with 1 mM DTT, pepstatin A, leupeptin, lysozyme, DNaseI, PMSF, and 0.1% Tween-20 and cell slurry was sonicated on ice. Lysed cells were centrifuged 20,000 xg for 30 minutes at 4C and soluble lysate was filtered prior to IMAC purification using 2×5mL His-Trap columns (GE Healthcare) affixed to an Akta Pure; elution buffer was 25 mM HEPES, pH 7.4, 150 mM NaCl, 500 mM imidazole. Fractions were pooled, additional protease inhibitors were added, and protein was dialyzed in 50 mM Bis-Tris, pH 6.7, 150 mM NaCl, 1 mM DTT with TEV protease to cleave the Trx-6xHis-tag at 4°C overnight. Cleaved protein was concentrated at 4°C using centrifugal concentrators prior to size exclusion chromatography using a Superdex 75 column. Protein was pooled and a second size exclusion chromatography step was performed using a Superdex 200 column using HEPES gel filtration buffer. Purity was confirmed to be ∼85% by SDS-PAGE and final protein concentration was confirmed with a combination of Nanodrop and Bradford Protein Assay.

## ACKNOWLEDGEMENTS

We thank Dr. Laura Solt for her contributions as a comentor to S.M. and Drs. Patrick Griffin, Mark Sundrud, and H. Jane Dyson for providing guidance to S.M. as her thesis committee. We are grateful to Dr. Albane le Maire (CNRS) for sharing the NCoR RID construct. This work was supported in part by National Institutes of Health (NIH) grants R01GM114420 (D.J.K.) and a Richard and Helen DeVos graduate fellowship award (S.M.). Use of the Stanford Synchrotron Radiation Lightsource, SLAC National Accelerator Laboratory, is supported by the U.S. Department of Energy, Office of Science, Office of Basic Energy Sciences under Contract No. DE-AC02-76SF00515. The SSRL Structural Molecular Biology Program is supported by the DOE Office of Biological and Environmental Research, and by the National Institutes of Health, National Institute of General Medical Sciences (including P41GM103393).

## AUTHOR CONTRIBUTIONS

S.M. and D.K. conceived and designed the research. S.M. expressed and purified protein and performed NMR analysis, biophysical, and biochemical assays. S.M., J.S., and P.M.T. collectively contributed to generating crystals, collecting and processing crystallography data, and solving the crystal structures. D.K. supervised the research and S.M. wrote the manuscript with input from all authors who approved the final version. The authors collectively declare no conflicts of interest in the completion of this study.

## DATA AVAILABILITY

Crystal structures of heme-bound REV-ERBβ LBD cobound to NCoR ID1 and ID2 motif peptides are available in the Protein Data Bank (PDB) under accession codes 6WMQ and 6WMS, respectively. NMR chemical shift assignments for heme-bound REV-ERBβ LBD are available in the Biological Magnetic Resonance Data Bank (BMRB) under ID 50251.

