## Supplementary Information for "Structural basis for heme-dependent NCoR binding to the transcriptional repressor REV-ERBβ"

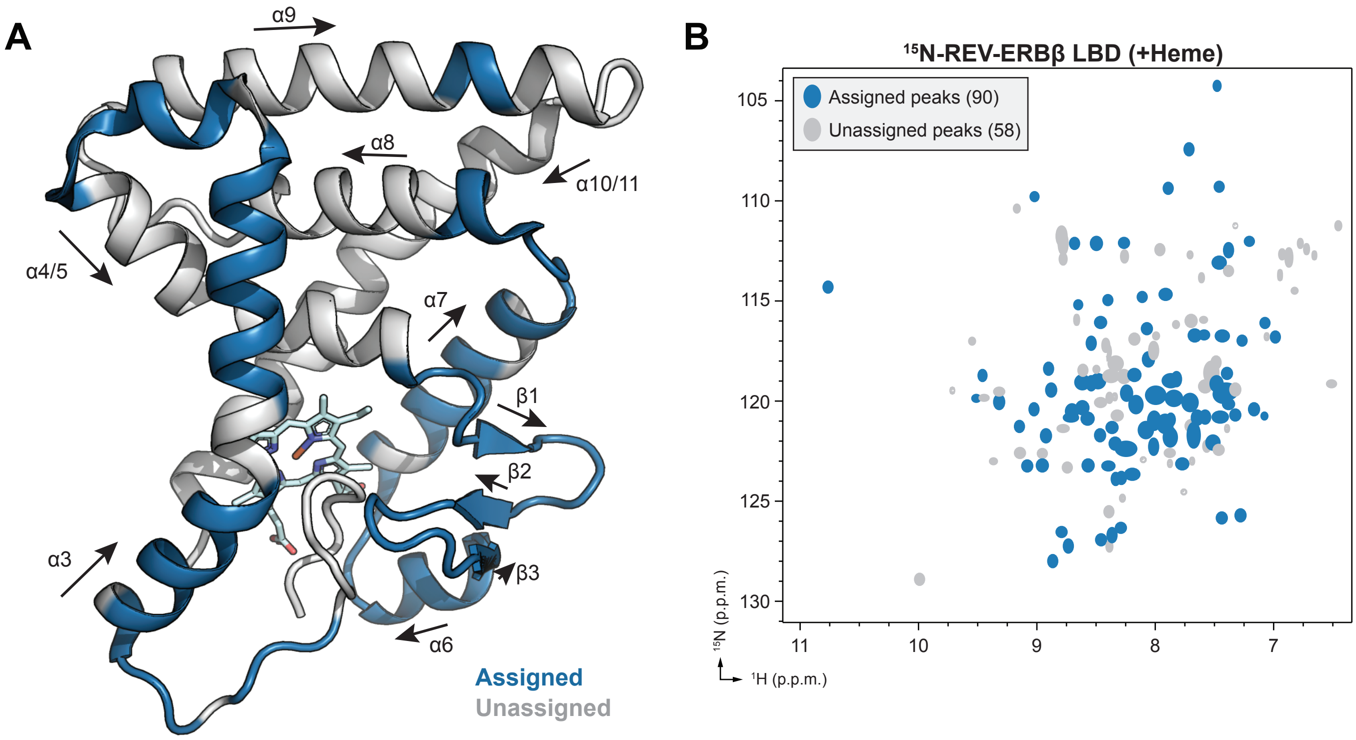

**Fig. S1. NMR chemical shift assignments of the heme-bound REV-ERBβ LBD spectrum.** (**A**) Residues with peak assignments in the 2D [$^{1}$H,$^{15}$N]-TROSY-HSQC spectrum of heme-bound $^{15}$N-REV-ERBβ LBD are colored dark blue on the heme-bound REV-ERBβ LBD structure (PDB 3CQV); unassigned residues are shown in gray; and secondary structure elements are marked with arrows. (**B**) 2D [$^{1}$H,$^{15}$N]-TROSY-HSQC spectrum of heme-bound REV-ERBβ LBD highlighting assigned (dark blue ellipses) and unassigned (gray ellipses) peaks.

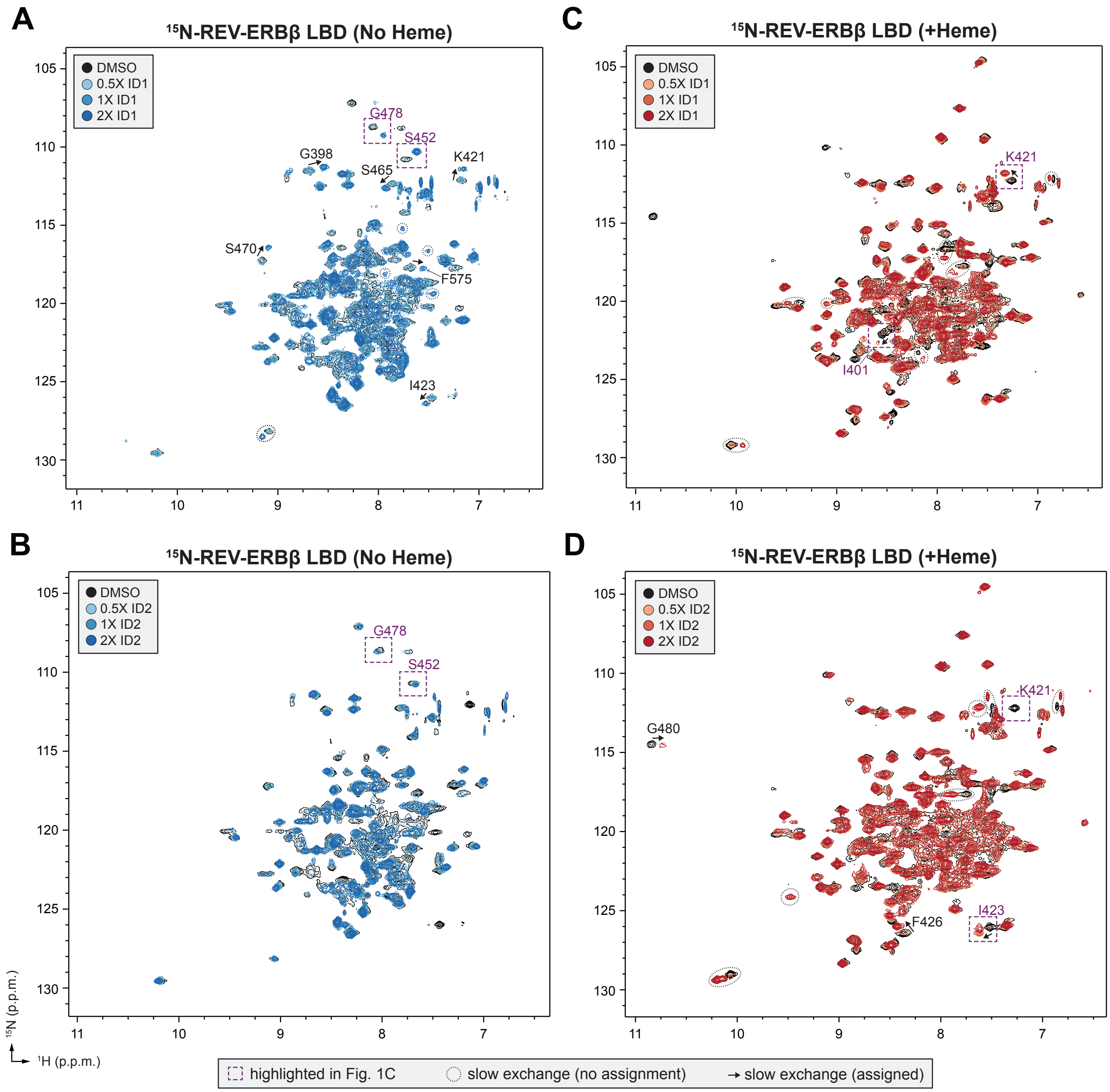

**Fig. S2. Full 2D [$^{1}$H,$^{15}$N]-TROSY-HSQC spectra of NCoR ID peptides titrated into REV-ERBβ LBD.** NCoR ID1 (**A**,**C**) or ID2 (**B**,**D**) peptide titrated into $^{15}$N-labeled REV-ERBβ LBD (vehicle is 0.8% DMSO) without (**A**,**B**) or with (**C**,**D**) heme added at the indicated peptide molar equivalents. Dashed purple boxes indicate the peaks highlighted in **Fig. 1C**. Black arrows indicate select peaks with slow exchange peak movements indicative of high affinity binding and are labeled with the corresponding residue. Dashed ovals indicate select peaks that move or appear in slow exchange, but were not assigned to an LBD residue.

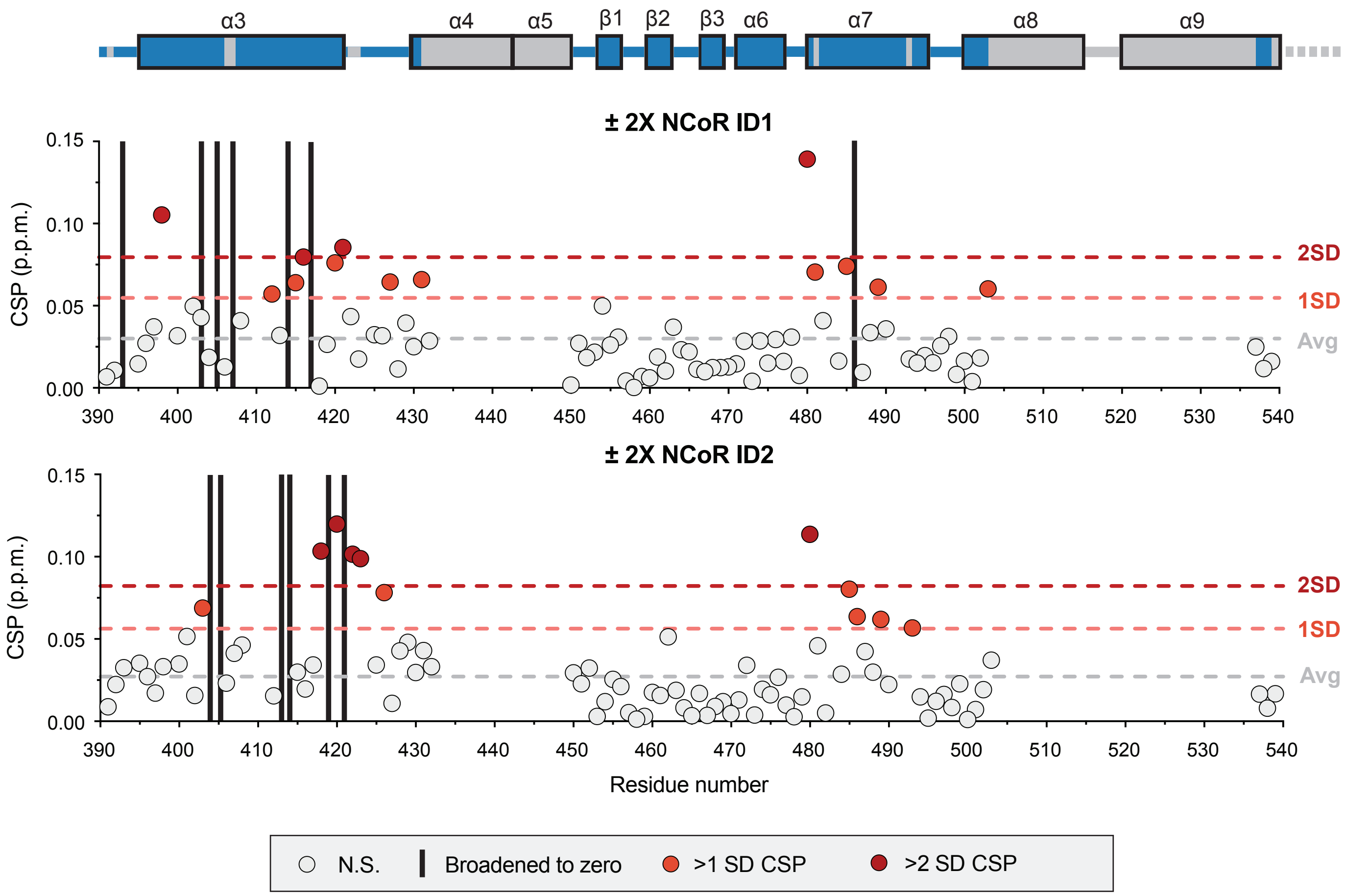

**Fig. S3. NMR structural footprinting analysis of NCoR ID peptides binding to heme-bound REV-ERBβ LBD.** Calculated chemical shift perturbations (CSPs) for assigned residues including notations for the average (Avg) CSP (gray dashed line) and CSPs more than 1 standard deviation (1SD; red dashed line) or 2 standard deviations (2SD; dark red dashed line) above the Avg CSP. REV-ERBβ secondary structures are shown at the top with boxes noting alpha helix (α) or beta-strand (β); dark blue regions have chemical shift assignments, and gray regions are unassigned.

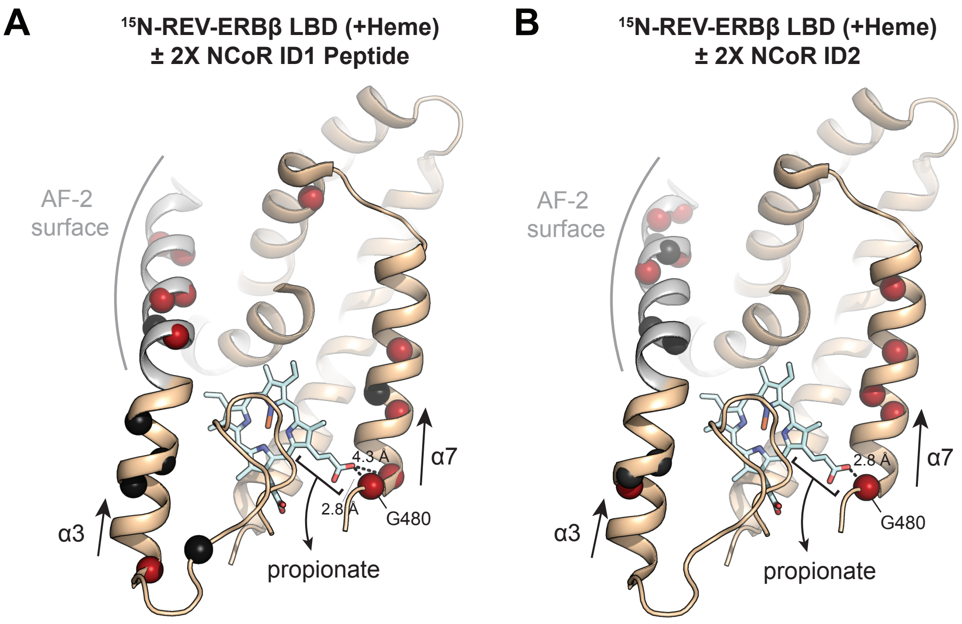

**Fig. S4. Putative mechanism for allosteric effects on helix 7 (α7) upon ID binding.** NMR structural footprinting effects mapped to the LBD (as in **Fig. 3D**) highlighting the effects on helix 7 that could be due to ID binding at the AF-2 surface allosterically shifting the heme propionate group, which hydrogen bonds to the backbone amide of G480 at the beginning of helix 7. The effects on helix 7 are observed for both $^{15}$N-REV-ERBβ LBD bound to heme in the presence of (**A**) ID1 peptide and (**B**) ID2 peptide.

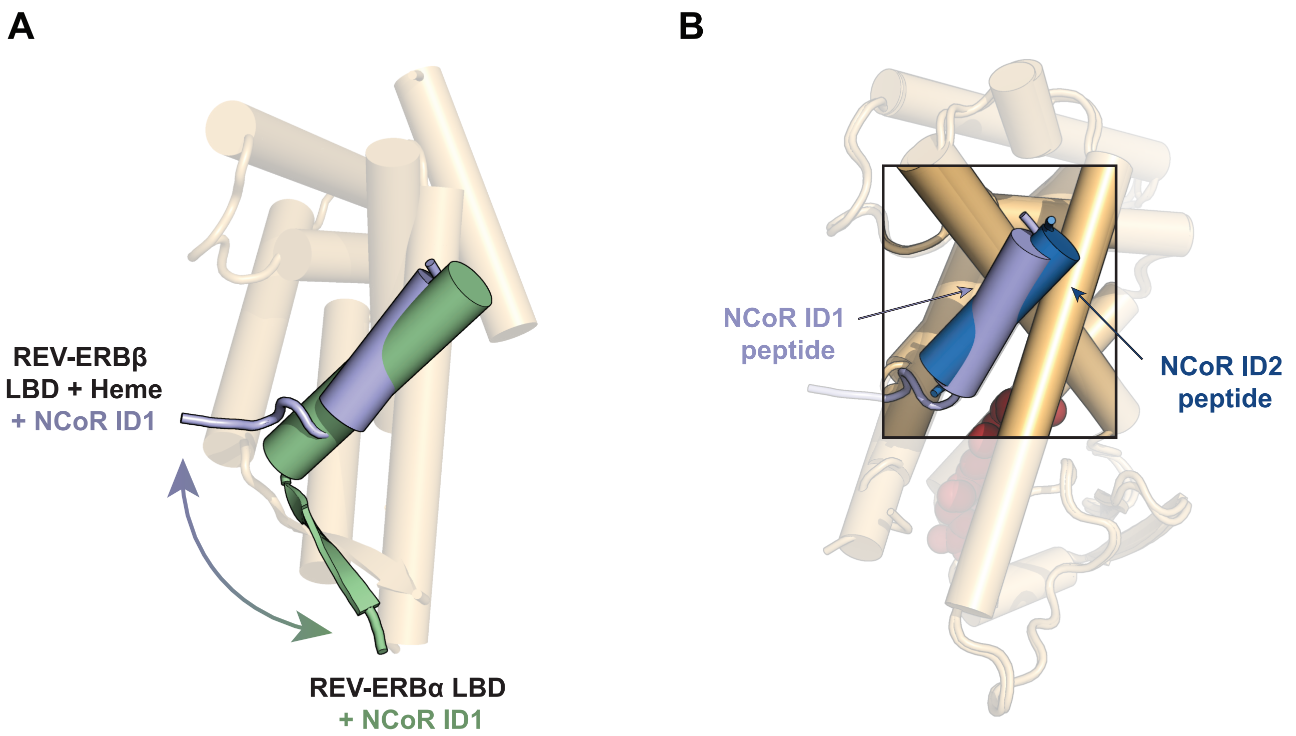

**Fig. S5. Disrupted beta-sheet and preserved alpha-helical interactions in the presence of heme.** (**A**) Overlay of the published apo-REV-ERBα LBD structure (light orange) bound to NCoR ID1 peptide (green) (PDB 3N00) and heme-bound REV-ERBβ LBD cobound NCoR ID1 peptide (purple) (PDB 6WMQ). α-helices are represented as cylinders. (**B**) Overlay of REV-ERBβ LBD structures (PDB 6WMQ and 6WMS) cobound to heme and NCoR ID peptides highlighting the similarity in the ID peptide α-helical binding mode.

**Table S1. Thermodynamic parameters of NCoR ID peptide binding to REV-ERBβ LBD.**

| | NCoR ID1 peptide | | NCoR ID2 peptide | |
|---|---|---|---|---|
| | REV-ERBβ (No Heme) | REV-ERBβ (+Heme) | REV-ERBβ (No Heme) | REV-ERBβ (+Heme) |
| [a]$K_D$ (μM) | 0.48<br>(95% [f]CI: 0.31 – 0.73) | 2.3<br>(95% CI: 1.3 – 4.2) | 16<br>(95% CI: 7.5 – 25) | 3.1<br>(95% CI: 2.3 – 4.4) |
| [b]ΔG (kcal/mol) | -8.6<br>(95% CI: -8.9 – -8.4) | -7.7<br>(95% CI: -8.0 – -7.3) | -6.5<br>(95% CI: -7.0 – -6.3) | -7.5<br>(95% CI: -7.7 – -7.3) |
| [c]ΔH (kcal/mol) | -7.5<br>(95% CI: -8.1 – -7.0) | 1.7<br>(95% CI: 1.5 – 1.9) | -3.3<br>(95% CI: -9.7 – -2.2) | 2.3<br>(95% CI: 2.1 – 2.4) |
| [d]TΔS (kcal/mol) | 1.1<br>(95% CI: 0.8 – 1.4) | 9.4<br>(95% CI: 9.2 – 9.5) | 3.2<br>(95% CI: -2.7 – 4.1) | 9.8<br>(95% CI: 9.7 – 9.8) |
| [e]N-value | 1 | 1 | 1 | 1 |

[a]Binding affinity.
[b]Free energy of binding.
[c]Enthalpy of binding.
[d]Temperature (T), which was constant at 25°C, and entropy (ΔS) of binding.
[e]Binding stoichiometry rounded to the nearest whole number.
[f]Confidence interval calculated using SEDPHAT analysis software.

**Table S2. X-ray crystallography data collection and refinement statistics.**

| | REV-ERBβ LBD cobound to heme and NCoR ID1 peptide | REV-ERBβ LBD cobound to heme and NCoR ID2 peptide |
|---|---|---|
| Data collection | | |
| Space group | C 1 2 1 | P 1 21 1 |
| Cell dimensions | | |
| a, b, c (Å) | 164.57, 48.50, 58.62 | 55.58, 73.40, 59.67 |
| α, β, γ (°) | 90, 105.3, 90 | 90, 101.91, 90 |
| Resolution | 46.38-2.55 (2.64-2.55) | 38.15-2.0 (2.07-2.0) |
| $R_{merge}$ | 0.095 (0.366) | 0.035 (0.313) |
| I / σ(I) | 6.83 (2.26) | 10.44 (1.94) |
| $CC_{1/2}$ in highest shell | 0.727 | 0.754 |
| Completeness (%) | 99.91 (100.00) | 99.09 (98.93) |
| Redundancy | 2.0 (2.0) | 2.0 (1.9) |
| Refinement | | |
| Resolution (Å) | 2.55 | 2.0 |
| No. of unique reflections | 14814 | 31527 |
| $R_{work}/R_{free}$ (%) | 20.6/27.4 | 18.8/23.9 |
| No. of atoms | | |
| Protein | 3369 | 3320 |
| Water | 58 | 204 |
| B-factors | | |
| Protein | 30.69 | 32.44 |
| Ligand | 19.04 | 23.13 |
| Water | 33.10 | 40.51 |
| Root mean square deviations | | |
| Bond lengths (Å) | 0.009 | 0.008 |
| Bond angles (°) | 1.19 | 0.90 |
| Ramachandran favored (%) | 95.42 | 98.77 |
| Ramachandran outliers (%) | 0.72 | 0.49 |
| PDB accession code | 6WMQ | 6WMS |

*Values in parentheses indicate highest resolution shell.

**Table S3. Thermodynamic parameters of NCoR RID binding to REV-ERBβ LBD.**

| | NCoR RID | |
|---|---|---|
| | REV-ERBβ (No Heme) | REV-ERBβ (+Heme) |
| [a]$K_D$ (μM) | 0.35<br>(95% [f]CI: 0.20 – 0.59) | 0.088<br>(95% CI: 0.036 – 0.19) |
| [b]ΔG (kcal/mol) | -8.8<br>(95% CI: -9.1 – -8.5) | -9.5<br>(95% CI: -10.2 – -9.2) |
| [c]ΔH (kcal/mol) | -10<br>(95% CI: -11 – -9.2) | 15<br>(95% CI: 13 – 16) |
| [d]TΔS (kcal/mol) | -1.2<br>(95% CI: -1.9 – -0.7) | 25<br>(95% CI: 23 – 25) |
| [e]N-value | 1 | 2 |

[a]Binding affinity.
[b]Free energy of binding.
[c]Enthalpy of binding.
[d]Temperature (T), which was constant at 25°C, and entropy (ΔS) of binding.
[e]Binding stoichiometry (REV-ERBβ LBD/NCoR RID) rounded to the nearest whole number.
[f]Confidence interval calculated using SEDPHAT analysis software.
